# Biomass allocation of trees in response to mono- and heterospecific neighbourhoods

**DOI:** 10.1101/2024.10.15.618432

**Authors:** Maria D. Perles-Garcia, Andreas Fichtner, Sylvia Haider, Hanjiao Gu, Wensheng Bu, Matthias Kunz, Werner Härdtle, Goddert von Oheimb

## Abstract

Carbon sequestration by trees is crucial to mitigate the effects of the current climate crisis. The extent to trees sequester and allocate carbon to above- or belowground structures in turn is mediated by neighbouring species. Although many studies have demonstrated positive effects of diverse neighbourhoods on a tree’s productivity, little is known about biomass allocation responses to mono-vs. heterospecific neighbourhoods. In the present study we quantified above- and belowground biomass production and root-to-shoot ratios (RSR) of trees grown in mono- and heterospecific neighbourhoods. To this end we analysed growth of mono- and heterospecific tree species pairs (TSPs) established in a greenhouse and a field experiment. In the greenhouse experiment response variables were measured after one year of growth after sapling harvest. In the field experiment, conducted in the context of a forest biodiversity experiment in subtropical China, we analysed biomass density and RSR over three years using terrestrial laser scanner and minirhizotrons. RSR of trees in heterospecific TSPs were significantly higher than in monospecific TSPs. In the greenhouse experiment, this was related to a stronger below-than aboveground overyielding in heterospecific TSPs. In the field experiment, trees in heterospecific TSPs showed a stronger increase in aboveground investments over time than in monospecific TSPs, indicating that positive diversity effects became stronger for aboveground structures with progressing tree development. Our findings are consistent with the optimal biomass partitioning theory and highlight the importance of tree-tree interactions on biomass allocation. Higher RSR in mixtures further suggest a higher resistance or resilience of tree saplings against environmental stressors related to climate change (drought, heat waves).

## 1. Introduction

Mitigating the effects of climate change is one of the major challenges of this century. Forests have the potential to fix CO_2_ from the atmosphere and to sequester carbon in the long term in their biomass. As a consequence, afforestation and reforestation measures are widely accepted as one of the most effective strategies to partly compensate for anthropogenic CO_2_ emissions (Bastin et al., 2019; Lewis et al., 2019). The acceleration of climate change in recent years has led to an increase in extreme events such as fires, droughts, floods or plagues (Kirilenko & Sedjo, 2007). Diversifying forests in turn may increase their resilience and their ability to fix carbon (C. L. C. Liu et al., 2018; Piotto, 2008). It is, therefore, important to understand the mechanisms through which the selection and combination of tree species may enhance forest growth and survival, and how these targets may be achieved by means of optimized afforestation strategies and forest restoration on devastated area.

In this context it is vital to know how trees may interact at the local neighbourhood level and in relation to neighbourhood species richness and composition (Fichtner et al., 2017). Several studies have demonstrated that tree growth and biomass allocation to different structures (roots, stems or branches) can vary depending on biotic and abiotic factors (Guillemot et al., 2020; Kunz et al., 2019; Williams et al., 2017). For example, competitive interactions between trees, but also species traits and trait plasticity as well as functional dissimilarity between neighbouring trees may mediate tree crown development and thus aboveground architecture (Guillemot et al., 2020; Hildebrand et al., 2020; Williams et al., 2021). In addition, tree-tree interactions may drive a tree’s biomass allocation within and between above- and belowground structures, which in turn affects a tree’s ability to compete for light and belowground resources such as nutrients and water (Beugnon et al., 2022; Kunz et al., 2019; Madsen et al., 2020). Even though local tree neighbourhoods have been shown to be one of the main drivers shaping a tree’s above- and belowground investments (Lang et al., 2010; Van de Peer et al., 2017; Yang et al., 2019), little is known about the underlying mechanisms and consequences for related traits such as root-to-shoot ratios. Root-to-shoot ratios (RSR), i.e. the ratio of the belowground biomass relative to the aboveground biomass, are measured from a tree individual or a forest stand and may have important implications for a tree’s or a stand’s ability to cope with environmental stressors such as drought events or heat waves (Qi et al., 2019). Modifying the neighbourhood species richness and composition of a focal tree thus might have consequences for its resistance or resilience to environmental stressors. Besides neighbourhood effects, RSR proved to be related to other factors such as latitude, stand age, nutrient availability, and climate (Qi et al., 2019). In addition, RSR are often used to estimate root biomass density from aboveground biomass, given that environmental factors are accounted for (Annighöfer et al., 2022; Cairns et al., 1997; Singnar et al., 2021).

Destructive methods allow direct measurements of above- and belowground biomass. In contrast, non-destructive methods may be less precise, but allow for continuous observations and longer-term measurements on shifts in allocation patterns in relation to the drivers of interest (e.g. tree-tree interactions). Non-destructive measures thus account for the dynamics of tree-tree interactions and time-related shifts in tree growth responses to biotic and abiotic environmental conditions (Li et al., 2014). In the present study we applied a combination of both approaches: In a greenhouse experiment tree seedlings were grown under controlled conditions and biomass of aboveground and belowground structures was quantified after sapling harvest one year after planting. In the field experiment we made use of the BEF-China experimental platform (Bruelheide et al., 2014) and quantified tree growth over three years using inventory and terrestrial laser scanning (TLS) data to quantify aboveground biomass development. In addition we installed minirhizotrons between each of the TSPs selected to examine changes in root biomass density (RBD) over time (Johnson et al., 2001)..

The main objective of the present study was to analyse how pairwise interactions between two adjacent, mono-vs. heterospecific trees species (i.e. tree species pairs, hereinafter abbreviated as TSPs) affect the trees’ biomass allocation and root-to-shoot ratios-. We hypothesize that (i) RSR would be higher in heterospecific TSPs because of relatively higher investments of saplings in belowground biomass, (ii) aboveground biomass investments might become more important over time in heterospecific TSPs, because positive diversity effects might be stronger aboveground with progressing crown development, and (iii) RSR are influenced by species composition and identity.

## 2. Material and methods

### 2.1. Greenhouse experiment

In autumn 2018, we collected seeds from eight broadleaved species in the Gutianshan National Nature Reserve (Zhejiang Province) which are also present in the biodiversity-ecosystem functioning experiment BEF China (see section 2.2). In spring 2019, seeds were germinated under controlled conditions in the greenhouse of the botanical garden from the MLU University Halle (Germany). From the individuals germinated 15 were selected by random from each species (8 species × 15 individuals = 120 individuals in total). In July 2019 the seedlings were measured (height above- and belowground, stem diameter, weight) and transplanted pairwise in 60 PVC tubes of 20 cm in diameter and 100 cm in length (i.e. two individuals per tube). The tubes were cut in two halves and re-assembled again so they could be opened without damaging the roots (Fig S1). We split the species into two sets of four species each and combine them in all possible combinations within the set. In total, 20 different species combinations were used: eight monospecific and 12 heterospecifics (Table S1).

All tubes in which both individuals survived were harvested in September 2020. In total, we analysed 98 trees, 40 and 58 of which grew in monospecific and heterospecific pairs, respectively. Table S1 shows the tree species combinations and the number of pairs that were sampled after one year for each treatment. For analyses, the root biomass was sampled by carefully cleaning roots from adhered soil material. Subsequently, aboveground and belowground biomass was dried at 80°C for 3 days and biomass dry weight of each tree quantified.

### 2.2. Field experiment (BEF China)

In addition to the one-year greenhouse experiment we used data from the BEF China field experiment. The BEF-China platform represents a Biodiversity–Ecosystem Functioning experiment situated in Jiangxi province (29.08°– 29.11°N, 117.90°– 117.93°E, 100– 300 m a.s.l.; Bruelheide et al., 2014). It has a subtropical climate, the mean temperature is 16,7°C and the mean precipitation is 1821 mm/year (Yang et al., 2013). Site A was planted in 2009. Each plot contains 400 trees planted in a regular grid with a distance between them of 1.29 m. Tree species richness of the plots ranges from monoculture to 24 species mix. TSPs can be monospecific pairs (i.e. both trees belong to the same tree species) or heterospecific pairs (i.e. the two trees belong to different tree species).

In this experiment we applied a non-destructive approach to study the growth behaviour of a total of 94 trees in 47 TSPs from 24 plots (Site A of the BEF-China experiment) over a three-year time period (2014-2016). All trees were identified by species and tree height and stem diameter at ground level were measured annually (between September and October). In addition, terrestrial laser scanning (TLS) was conducted each year in March on 14 of the 24 plots (see description below). Table S2 shows the species combinations, the number of TSPs analysed each year and the number of plots in which each species combination was present. In May 2014 minirhizotrons were installed in the middle of the two trees. From the minirhizotron tube, photos were taken annually to document the trees’ rooting pattern (see description below).

#### 2.2.1. Aboveground biomass estimation

TLS was conducted under leaf-off conditions, using the laser scanner FARO Focus S120. The TLS campaigns included 714 trees. We scanned a total of 18 TSPs out of the 47 TSPs included in this study. We generated quantitative structure models (QSMs) from the point clouds to quantify the wood volume of individual trees. Co-registration and point cloud segmentation were carried out manually using FARO Scene software (V. 5.2.6), RiSCAN PRO (v2.6.2) and Bentley Pointools (v1.5 Pro). For QSMs, we used the TreeQSM software (Åkerblom, 2017; see Kunz et al. 2019 for more details)

To estimate the volume of TSPs that were not scanned and in order to have a consistent data set, we used the random forest algorithm. We used height, stem diameter at ground level and species as explanatory variables and trained the algorithm with a subset of 665 scanned multiple years, n = 1326, with the volume derived from TLS as a response. We performed the analysis using R 4.1. and the randomForest package (Liaw & Wiener, 2002). We validated the results with a subsample of 427 trees scanned multiple years, n = 591 (RMSE = 5.04, MAE = 2.58, R^2^ = 0.79), and with the 36 trees from the 18 TSPs included in the study (RMSE = 8.30, MAE = 6.64, R^2^ = 0.77).

Finally, we used the modelled volume and the wood density of each tree species (taken from Kröber et al., 2014) to calculate the aboveground biomass of each tree.

#### 2.2.2. Belowground biomass estimation

The minirhizotron tubes were installed at a 30° angle to a soil depth of 30 cm (diagonal length of 60 cm). Each minirhizotron was installed in the middle of two neighbouring trees. Before installing the minirhizotrons, we removed understory plants and leaf litter from the ground. Plugs were used to seal the bottom of all pipes, and plastic covers and self-sealing bags were used to seal the top (Fig S2). The sealings were applied to prevent contamination with organic matter and the infiltration of water. Black tapes followed by white tapes were applied to the aboveground part of each tube in order to prevent heat absorption caused by penetrating light (Kou et al., 2018). To minimize soil disturbance, leaf litter was used to cover the ground around the tubes. Root scanning started seven months after installation of the minirhizotrons, to allow stabilization of the soil (Hansson et al., 2013).

To improve fine root turnover analyses, we acquired coloured root images (18 mm × 14 mm) at the same soil depth in November 2014, May 2015 and November 2016. BTC-100 camera system (Bartz Technology, Santa Barbara, Calif.) was used for root images and to document root growth. We collected about 160 images in four directions in each tube. WinRhizo Tron MF (Regent Software, Canada) was used to process the images and to get data on total root length and alive root length. The identification of living and dead roots followed the definition of Wells & Eissenstat (2001), according to which living roots show a white and brown colour, while dead roots exhibit an exfoliated cortex, wrinkled epidermis, or black colour.

We quantified the following belowground traits: specific root length (SRL), root length density per unit volume (RLD_v_), and root biomass density per unit volume (RBD_v_). RLD is an important indicator reflecting the below-ground competitiveness of trees. The calculation of RLD_V_ is as follows:

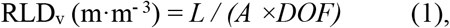

where *L* is the root length inferred from the minirhizotron images (m), *A* is an area of minirhizotron image observed, *DOF* is field depth (m), that was set to 0.002 m (Steele et al., 1997).

For the calculation of SRL, intact fine roots were excavated from the top 30 cm of undisturbed soil through drilled soil samples in each plot and transported to the laboratory in a cooler box with ice packs in September 2015. For each plot, at least 6 individuals per species were sampled in plots with tree species richness of one and 3 individuals per species were sampled in plots with tree species richness greater than one and heterospecific direct neighbours. After gently cleaning the soil and organic particles of the fine roots with tap water, the fine roots were divided into 5 functional groups according to Pregitzer et al. (2002). These five groups were split into two types; that is, the absorptive fine roots (orders 1-3) and transport fine roots (orders 4, and 5). Then, roots of each functional type were scanned by an Epson scanner and analyzed by WinRHIZO (Regent Software, Canada) to get data on total root length and average diameter. After scanning, these samples were dried in an oven at 60°C for over 48 h until they became constant weight and weighed. The calculation of SRL of each order is as follows:

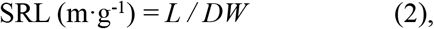

where *L* is the root length observed in each scanned image (m), *DW* is the root dry weight (g).

Since minirhizotrons were installed in the middle of two neighbour individuals, the SRL in mixtures represents the mean specific root length of absorptive fine roots of the two species. RBD was estimated in combination with SRL data obtained by soil drilling (Shi et al., 2006). The calculation of RBD_v_ is as follows:

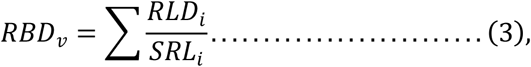

where *RBD*_*v*_ is the root biomass density per unit volume (g·m^-3^). *RLD*_*i*_ is the *RLD*_*v*_ (m·m^-3^) of diameter *i*; and *SRL*_*i*_ is the *SRL* (m·g^-1^) of diameter i.

### 2.3. Statistical analysis

We applied linear mixed-effects models to examine how pairwise tree interactions affect above- and belowground biomass, and RSR, and whether this effect varies over time. Specifically, we analysed the changes depending on whether the trees grew in mono- or heterospecific pairs. We used different approaches:

In the greenhouse experiment, we used the measured dry weight for above- and belowground biomass, and calculated the RSR by dividing the root dry weight of each individual by the dry weight of its shoots. We used TSP species richness (mono- or heterospecific) as predictor and species and neighbour species as random intercepts. We also consider the result of the TSPs as a whole, calculating the sum of the biomass of both trees. In this case, the combination of species was used as a random intercept. The model was fitted separately to predict aboveground biomass weight, belowground biomass weight, and RSR.

In the case of the field experiment, we analysed the response of the TSP: We divided the RBD by the shoot biomass density (SBD), calculated as the biomass (g) divided by two times the distance between the trees and the mean height of the studied trees (2.58 m x 2.58 m x 3.40 m). We fitted different linear mixed-effects models using the RBD, SBD and RSR of the TSP as the response variable and we tested the effect of TSP diversity (mono- or heterospecific), year and the interaction between both. Year 0 was considered the year of the planting of Site A (i.e. 2009). Species combination of a specific TSP and study plot were used as random intercepts. To assure model assumptions (normality, homogeneity and independence; Zuur et al., 2009), we square root transform the dependent variables in both data sets (greenhouse and field experiment data).

All statistical analyses were performed in R 4.0.2 (R Core Team, 2018) using the packages lme4 (Bates et al., 2015), lmerTest (Kuznetsova et al., 2017) and sjPlot (Lüdecke, 2020).

## 3. Results

Overall, we observed a significant relation between species richnes of TSPs and our three response variables (above- and belowground biomass (or biomass density), and RSR; Table 1, Table 2, Fig 1, Fig 2). One year after planting, seedlings from the greenhouse experiment had significantly more biomass above- and belowground in mixtures than in monocultures (Table 1, Fig 1). In addition, RSR was significantly higher in mixtures (Table 1, Fig 1). However, when we analyse this data grouped by TSPs, the p-value was not significant for any of the three models (Table S3).

**Table 1.** Results of mixed-effects models for the effects of pairwise species richness (mono- or heterospecific) on the root to shoot ratio (RSR, squared root transformed), aboveground dry biomass (AG_weight, squared root transform), and belowground dry biomass (BG_weight, squared root transform) for 98 trees planted under control conditions in the greenhouse experiment.

| Fixed effect | sqrt-RSR |  |  |  | sqrt-AG_weight (g) |  |  |  | sqrt-BG_weight (g) |  |  |  |
| --- | --- | --- | --- | --- | --- | --- | --- | --- | --- | --- | --- | --- |
|  | df<sub>num</sub> | df<sub>den</sub> | F | p | df<sub>num</sub> | df<sub>den</sub> | F | p | df<sub>num</sub> | df<sub>den</sub> | F | p |
| Intercept | - | - | - | < 0.001 | - | - | - | < 0.05 | - | - | - | < 0.05 |
| Species richness | 1 | 83.391 | 5.311 | < 0.05 | 1 | 83.109 | 8.057 | < 0.01 | 1 | 83.099 | 15.389 | < 0.001 |
| Marginal R<sup>2</sup> | 0.027 |  |  |  | 0.011 |  |  |  | 0.018 |  |  |  |
| Conditional R<sup>2</sup> | 0.512 |  |  |  | 0.868 |  |  |  | 0.891 |  |  |  |

dfnum, numerator degrees of freedom; dfden, denominator degrees of freedom.

**Table 2.** Results of mixed-effects models for the effects of pairwise diversity (mono- or hetero- specific, “Spec_Div”), year and its interaction on the root to shoot ratio (RSR, squared root transformed), shoot biomass density (SBD, squared root transform), and root biomass density (RBD, squared root transform). The model was fitted to 47 TSPs measured during three years (2014, 2015, and 2016), in a total of 126 observations.

| Fixed effect | sqrt-RSR |  |  |  | sqrt-SBD (g/m<sup>3</sup>) |  |  |  | sqrt-RBD (g/m<sup>3</sup>) |  |  |  |
| --- | --- | --- | --- | --- | --- | --- | --- | --- | --- | --- | --- | --- |
|  | df<sub>num</sub> | df<sub>den</sub> | F | p | df<sub>num</sub> | df<sub>den</sub> | F | p | df<sub>num</sub> | df<sub>den</sub> | F | p |
| Intercept | - | - | - | < 0.01 | - | - | - | < 0.001 | - | - | - | < 0.001 |
| Species richness | 1 | 111.188 | 14.594 | < 0.001 | 1 | 120.811 | 8.222 | < 0.01 | 1 | 85.684 | 2.7494 | 0.101 |
| Year | 1 | 81.228 | 44.481 | < 0.001 | 1 | 80.177 | 40.218 | < 0.05 | 1 | 77.311 | 98.606 | < 0.001 |
| Species richness×Year | 1 | 81.144 | 12.927 | < 0.001 | 1 | 80.138 | 9.146 | < 0.01 | 1 | 77.33 | 2.277 | 0.135 |
| Marginal R<sup>2</sup> | 0.097 |  |  |  | 0.061 |  |  |  | 0.373 |  |  |  |
| Conditional R<sup>2</sup> | 0.821 |  |  |  | 0.922 |  |  |  | 0.576 |  |  |  |

dfnum, numerator degrees of freedom; dfden, denominator degrees of freedom.

**Figure 1.**
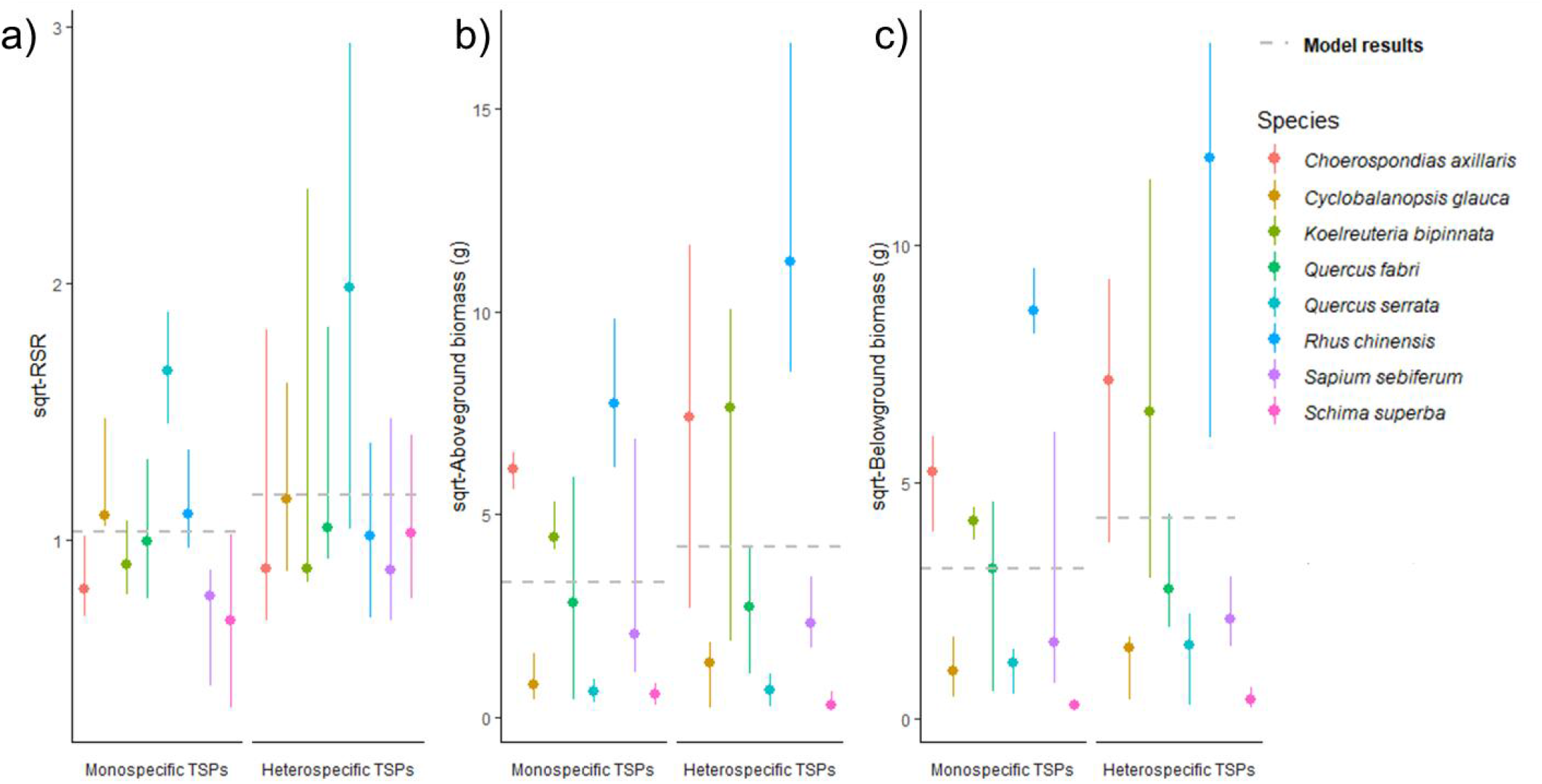
Range bar of observed values and model results for the three response variables measured in the greenhouse: (a) shows root-to-shoot ratio (RSR, squared root transformed), (b) aboveground dry biomass squared root transform, and (c) belowground dry biomass squared root transform. Dots represent the median of the observed values grouped by species and by pairwise diversity. Solid lines represent the range of the results. The grey dashed line represents the predicted values of the models.

**Figure 2.**
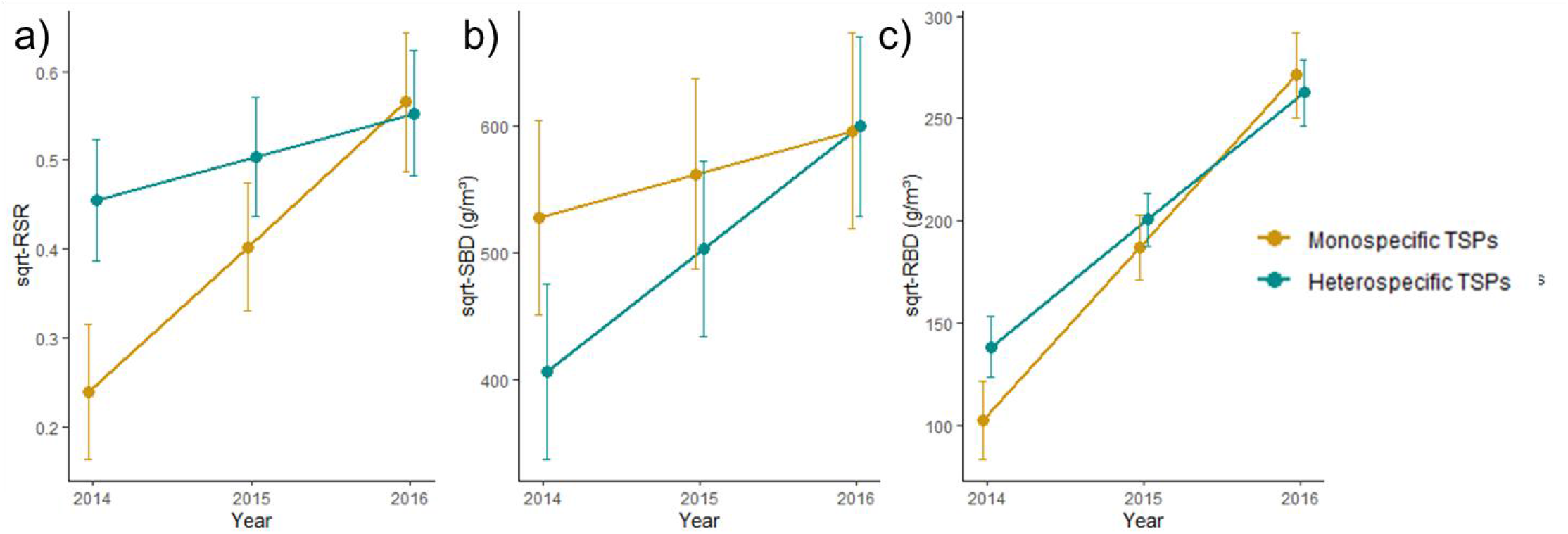
Effects of pairwise species diversity (mono- or hetero- specific pairs) on (a) root to shoot ratio (RSR, squared root transformed), (b) shoot biomass density (SBD, squared root transform), and (c) root biomass density (RBD, squared root transform). Lines represent mixed-effects model fits, dots represent the mean predicted value for each year, and the corresponding standard errors are represented as error bars.

We found comparable results in the field experiment in that seedlings showed higher RSR and higher values for SBD in hetero-than in monospecific TSPs, but differences for RBD were not significant (Table 2). In addition, tree growth responses strengthened over time, indicated by a positive and significant “Year”-effect across response variables (p < 0.05 for all three variables; Table 2). However, time effects on RSR and SBD were different for mono-und heterospecific TSPs, indicated by the significant two-way interaction between “Year” and “Species richness”. As shown in Fig 2, trees in heterospecific TSPs showed a stronger increase in aboveground biomass investments over time than in monospecific TSPs. As a result, the increase in RSR over time was stronger in monospecific TSPs (Fig 2a). In contrast, the “Year × Species richness” interaction was not significant for RBD (p = 0.135, Table 2).

Heterospecific TSPs benefit both, above- and belowground over time, with effects being higher for RBD: In 2016 the estimate RBD was 90% higher than in 2014, compared to a 48% increase in SBD. Meanwhile, we estimated a much more pronounced increase of RBD in monospecific TSPs: While its estimated increase in SBD between 2014 and 2016 was 13%, in RBD was 165%

## 4. Discussion

There is evidence that neighbourhood interactions can modulate the aboveground tree architecture as a result of interspecific differences in morphological and physiological tree traits (Hildebrand et al., 2020; Perles-Garcia et al., 2022; Williams et al., 2017). However, due to sampling difficulties few studies have analysed above- and belowground biomass allocation patterns of trees in relation to con- and heterospecific neighbourhoods. Instead, many studies focused on tree growth responses and crown formation to abiotic factors such as nutrient availability or climate (Y. Liu et al., 2014; Meier & Leuschner, 2008; Trubat et al., 2011; Wright, 2019), or compared estimations of RSR in different biomes (Ma et al., 2021; Mokany et al., 2006; Qi et al., 2019). To address this gap in knowledge, the present study quantified how biomass allocation of different tree species responds to con- and heterospecific neighbourhoods, using two different experimental approaches. Our results evidenced that tree-tree interactions may have significant effects on the biomass allocation of tree individuals, with possible consequences for growth responses to and performance under environmental stressors.

In the greenhouse experiment, trees in heterospecific TSPs showed a higher above- and belowground productivity than trees in monospecific TSPs. This finding is in agreement with previous studies, according to which more diverse forests show overyielding compared to monocultures, mainly attributable to complementarity between different tree species (Huang et al., 2018; Liang et al., 2016; Paquette & Messier, 2011; van der Plas, 2019). However, this finding became non-significant when comparing TSPs instead of tree individuals, likely indicating a strong effect of the species composition of the respective TSPs. This interpretation is supported by the finding that all models showed distinct differences between R^2^_marginal_ and R^2^_conditional_ (see Tables 1 and 2 and further discussion below).

As hypothesized, RSR were higher for saplings in heterospecific than in monospecific TSPs (for both experiments). Since survival of tree seedlings often depends on a rapid development of their root system to ensure a sufficient supply with belowground resources (Grossnickle, 2012), competitive interactions initially might have been stronger belowground, particularly in the greenhouse experiment with limited roots space within the root tubes. This would explain why positive diversity effects for saplings might be stronger belowground, leading to higher biomass production in heterospecific TSPs. Our interpretation is supported by the finding that overyielding was stronger for belowground than for aboveground biomass in the greenhouse experiment, probably because of an optimised (complementary) use of resources due to belowground niche differentiation. Our findings are in agreement with those from Ravenek et al. (2014), who also observed overyielding in root biomass with increasing species richness.

In the field experiment, RSR showed a steady increase over time for trees in both mono- and heterospecific TSPs, attributable to a stronger increase in belowground investments. This plasticity in allocation can be interpreted as a general change in a tree’s allometric trajectory with increasing tree age and thus, as allometric growth (“apparent plasticity”) in the sense of Weiner (2004). However, as indicated by a significant “Year × Species richness” interaction, trees in monospecific TSPs showed a significantly stronger increase in their RSR than trees in heterospecific TSPs. Since TSP-related differences in the increase of RBD were non-significant, the weaker increase in RSR observed in heterospecific TSPs is solely caused by a stronger increase in aboveground biomass investments over time. This confirms our second hypothesis that aboveground biomass investments might become more important over time in heterospecific TSPs, because positive diversity effects (e.g. because of niche partitioning and complementary resource use) might be stronger aboveground with progressing crown development (Jucker et al., 2020). As a result, trees grown in heterospecific neighbourhoods may increase their aboveground investments, concomitantly optimize resource acquisition (e.g. light foraging) and minimize interspecific competition. In contrast to heterospecific TSPs and in accordance with the optimal biomass partitioning theory (Bloom & Mooney, 1985), monospecific pairs had higher aboveground investments in 2014, specifically in the elongation of the primary stem, to avoid shading by competing neighbours with similar growth rates. Due to an adjustment of biomass allocation to the most limiting resource (Bloom & Mooney, 1985), trees in monospecific pairs showed an increasing (relative) investment to belowground structures. However, further studies covering longer-time measurement series are required to determine whether this trend persists over time.

Our result that mixtures show lower aboveground biomass values in the early years of development is in agreement with findings from Jucker et al. (2020), but contradicts the findings of Huang et al. (2018), who observed consistent higher productivity at the plot level with increasing tree species richness in the BEF-China experiment for the years 2013-2017. These differences could be due to the fact that we focused on the TSPs between which the minirhizotrons were installed, and we studied the years 2014-2016. Also, SBD accounts for the sum of the biomasses of both trees of the TSP, which could mean that fast-growing species will have lower values when mixed with slow-growing species.

We found that variation in above, below and RSR is largely explained by species identity effects (i.e. species combination of a given TSP) rather than species richness effects (mono-versus heterospecific TSP), which supports our third hypothesis (also see Fig S3 illustrating the biomass of tree individuals from BEF China by species). We assume that the lower values in aboveground biomass in the fifth and sixth year after planting are related to species-specific traits such slow or fast-growing strategies, but also to the fact that other neighbouring trees surrounding the TSPs were not considered (i.e. four of the monospecific TSPs studied were not grown in plots planted as monocultures; this concerns TSPs of *Sapindus mukorossi, Castanopsis sclerophylla, Cyclobalanopsis glauca*, and *Cyclobalanopsis myrsinifolia*, where one of the TSPs considered as monospecific grew in a plot with tree species richness of two, four, four and four respectively).

In conclusion our findings demonstrate that tree-tree interactions at the neighbourhood level may significantly affect allometric growth of trees in terms of above-and belowground biomass investments, which in turn might influence a tree’s growth vigour and performance under shifting environmental conditions. Higher RSR in mixtures suggest a higher resistance or resilience of tree saplings against environmental stressors related to climate change (drought, heat waves). The establishment of diverse tree plantations in the context of afforestation or restoration measures thus might contribute to more stable tree communities under changing climatic conditions.

