## Supplementary information for "Biomass allocation of trees in response to mono- and heterospecific neighbourhoods"

Differences in above- and belowground biomass allocation between mono- and heterospecific tree species pairs

#### Table of contents

##### *Supporting Tables*


- **Table S1**. Tree species combinations measured in the greenhouse
- **Table S2.** Tree species combinations measured in BEF China
- **Table S3.** Mix-effect model results for the greenhouse data grouped in TSPs


##### *Supporting Figures*


- **Fig. S1.** Boxplots of estimated biomass of the trees from BEF China by species

### Supporting tables

*Table S1. Tree species combinations and the number of pairs that were sampled from the greenhouse experiment.*

| | | Number |
|---|---|---|
| *Choerospondias axillaris_Choerospondias axillaris* | mono | 3 |
| *Cyclobalanopsis glauca_Cyclobalanopsis glauca* | mono | 3 |
| *Koelreuteria bipinnata_Koelreuteria bipinnata* | mono | 3 |
| *Quercus fabri_Quercus fabri* | mono | 3 |
| *Quercus serrata_Quercus serrata* | mono | 2 |
| *Rhus chinensis_Rhus chinensis* | mono | 2 |
| *Sapium sebiferum_Sapium sebiferum* | mono | 3 |
| *Schima superba_Schima superba* | mono | 1 |
| *Choerospondias axillaris_Koelreuteria bipinnata* | mix | 3 |
| *Choerospondias axillaris_Quercus serrata* | mix | 1 |
| *Choerospondias axillaris_Sapium sebiferum* | mix | 3 |
| *Cyclobalanopsis glauca_Quercus fabri* | mix | 3 |
| *Cyclobalanopsis glauca_Rhus chinensis* | mix | 3 |
| *Cyclobalanopsis glauca_Schima superba* | mix | 2 |
| *Koelreuteria bipinnata_Quercus serrata* | mix | 2 |
| *Koelreuteria bipinnata_Sapium sebiferum* | mix | 3 |
| *Quercus fabri_Rhus chinensis* | mix | 3 |
| *Quercus fabri_Schima superba* | mix | 2 |
| *Quercus serrata_Sapium sebiferum* | mix | 1 |
| *Rhus chinensis_Schima superba* | mix | 3 |

*Table S2. Species combinations, number of TSPs analysed each year, and number of plots from BEF China in which each species combination was present.*

| | | 2014 | 2015 | 2016 | Plots |
|---|---|---|---|---|---|
| *Castanea henryi_Castanea henryi* | mono | 1 | 0 | 0 | 1 |
| *Castanopsis sclerophylla_Castanopsis sclerophylla* | mono | 2 | 2 | 2 | 2 |
| *Choerospondias axillaris_Choerospondias axillaris* | mono | 1 | 1 | 0 | 1 |
| *Cyclobalanopsis glauca_Cyclobalanopsis glauca* | mono | 2 | 2 | 2 | 2 |
| *Cyclobalanopsis myrsinifolia_Cyclobalanopsis myrsinifolia* | mono | 1 | 1 | 1 | 1 |
| *Liquidambar formosana_Liquidambar formosana* | mono | 1 | 1 | 1 | 1 |
| *Nyssa sinensis_Nyssa sinensis* | mono | 1 | 1 | 1 | 1 |
| *Quercus fabri_Quercus fabri* | mono | 1 | 1 | 1 | 1 |
| *Quercus serrata_Quercus serrata* | mono | 1 | 1 | 0 | 1 |
| *Sapindus mukorossi_Sapindus mukorossi* | mono | 2 | 2 | 2 | 2 |
| *Sapium sebiferum_Sapium sebiferum* | mono | 1 | 1 | 1 | 1 |
| *Castanea henryi_Diospyros glaucifolia* | mix | 1 | 1 | 1 | 1 |
| *Castanea henryi_Koelreuteria bipinnata* | mix | 1 | 1 | 0 | 1 |
| *Castanea henryi_Nyssa sinensis* | mix | 1 | 1 | 1 | 1 |
| *Castanopsis sclerophylla_Choerospondias axillaris* | mix | 1 | 1 | 0 | 1 |
| *Castanopsis sclerophylla_Cyclobalanopsis myrsinifolia* | mix | 1 | 1 | 1 | 1 |
| *Castanopsis sclerophylla_Quercus serrata* | mix | 2 | 2 | 1 | 2 |
| *Choerospondias axillaris_Lithocarpus glaber* | mix | 1 | 1 | 1 | 1 |

| | | | | | |
|---|---|---|---|---|---|
| *Choerospondias axillaris_Quercus serrata* | mix | 1 | 1 | 1 | 1 |
| *Choerospondias axillaris_Sapium sebiferum* | mix | 2 | 2 | 2 | 2 |
| *Cyclobalanopsis glauca_Lithocarpus glaber* | mix | 1 | 1 | 1 | 1 |
| *Cyclobalanopsis glauca_Nyssa sinensis* | mix | 1 | 1 | 1 | 1 |
| *Cyclobalanopsis glauca_Quercus fabri* | mix | 1 | 1 | 1 | 1 |
| *Cyclobalanopsis glauca_Schima superba* | mix | 1 | 1 | 1 | 1 |
| *Cyclobalanopsis myrsinifolia_Lithocarpus glaber* | mix | 1 | 1 | 1 | 1 |
| *Cyclobalanopsis myrsinifolia_Schima superba* | mix | 1 | 1 | 0 | 1 |
| *Koelreuteria bipinnata_Lithocarpus glaber* | mix | 3 | 3 | 3 | 2 |
| *Liquidambar formosana_Lithocarpus glaber* | mix | 1 | 1 | 1 | 1 |
| *Liquidambar formosana_Quercus serrata* | mix | 1 | 1 | 1 | 1 |
| *Liquidambar formosana_Sapindus mukorossi* | mix | 4 | 4 | 4 | 2 |
| *Liquidambar formosana_Sapium sebiferum* | mix | 1 | 1 | 1 | 1 |
| *Lithocarpus glaber_Quercus fabri* | mix | 1 | 1 | 0 | 1 |
| *Lithocarpus glaber_Rhus chinensis* | mix | 1 | 0 | 0 | 1 |
| *Quercus fabri_Sapium sebiferum* | mix | 1 | 1 | 0 | 1 |
| *Quercus fabri_Schima superba* | mix | 1 | 0 | 0 | 1 |
| *Quercus serrata_Schima superba* | mix | 1 | 1 | 1 | 1 |
| *Rhus chinensis_Schima superba* | mix | 1 | 1 | 0 | 1 |
| | **Total:** | **47** | **44** | **35** | **24** |

*Table S3. Results of mixed-effects models for the effects of pairwise diversity (mono- or hetero- specific, "Spec_Div") on the root to shoot ratio (RSR, squared root transformed), aboveground dry biomass (AG_weight, squared root transform), and belowground dry biomass (BG_weight, squared root transform) for 49 TSPs planted under control conditions in a greenhouse experiment.*

| | **sqrt-RSR** | | | | **sqrt-AG_weight (g)** | | | | **sqrt-BG_weight (g)** | | | |
|---|---|---|---|---|---|---|---|---|---|---|---|---|
| Fixed effect | $df_{num}$ | $df_{den}$ | F | p | $df_{num}$ | $df_{den}$ | F | p | $df_{num}$ | $df_{den}$ | F | p |
| Intercept | - | - | - | < 0.001 | - | - | - | < 0.001 | - | - | - | < 0.001 |
| Spec_Div | 1 | 10.249 | 0.8272 | 0.384 | 1 | 17.795 | 2.2266 | 0.1532 | 1 | 17.783 | 2.4179 | 0.1376 |
| Marginal $R^2$ | | | | 0.026 | | | | 0.092 | | | | 0.101 |
| Conditional $R^2$ | | | | 0.352 | | | | 0.810 | | | | 0.837 |

dfnum, numerator degrees of freedom; dfden, denominator degrees of freedom.

### Supporting figures

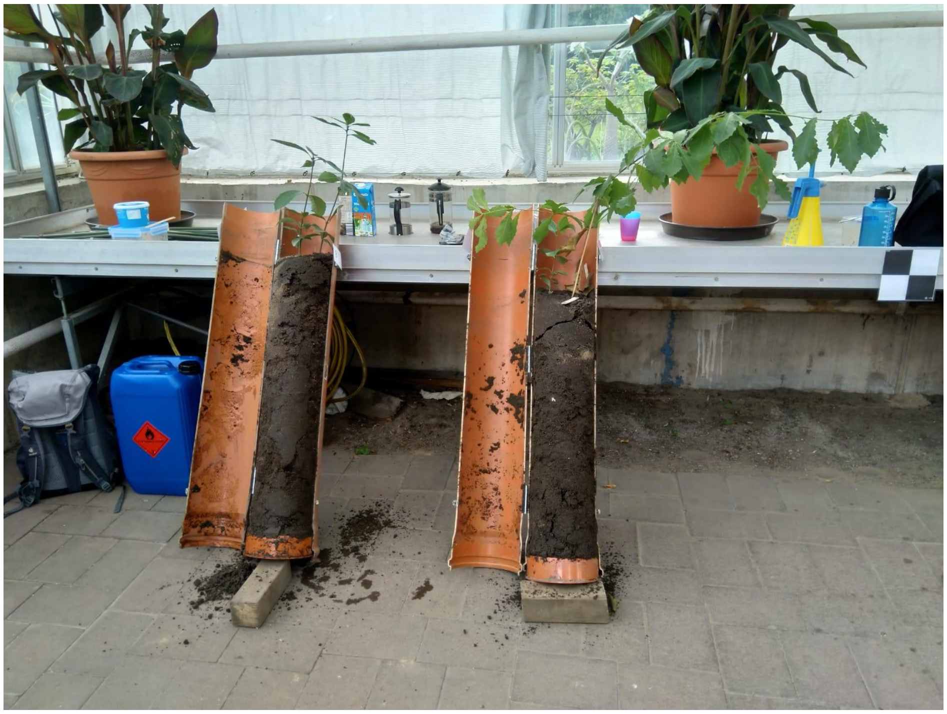

**Figure S1.** Photograph of the greenhouse experiment: two TSPs in their respective tubes just prior to harvesting (September 2020).

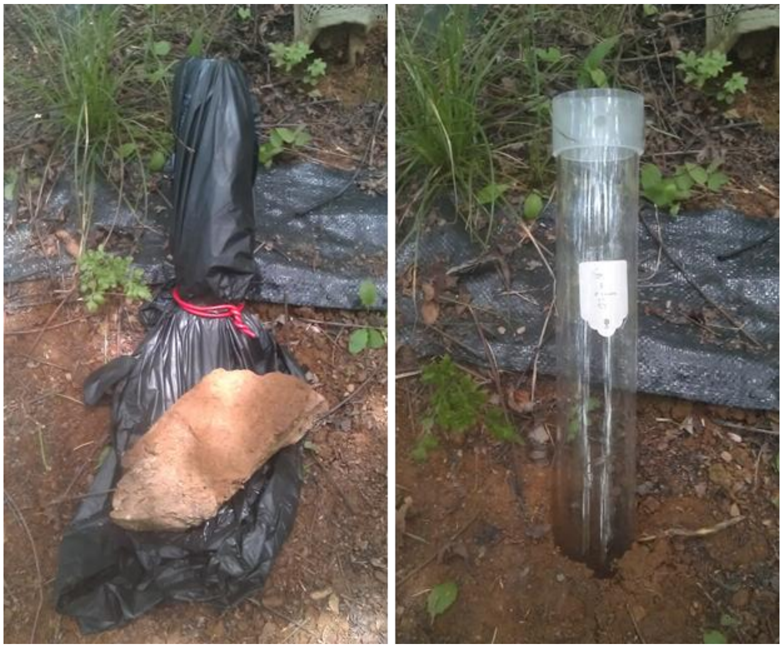

**Figure S2.** Photographs of one minirhizotron installed in BEF China with a plastic cover (left) and unsealed (right).

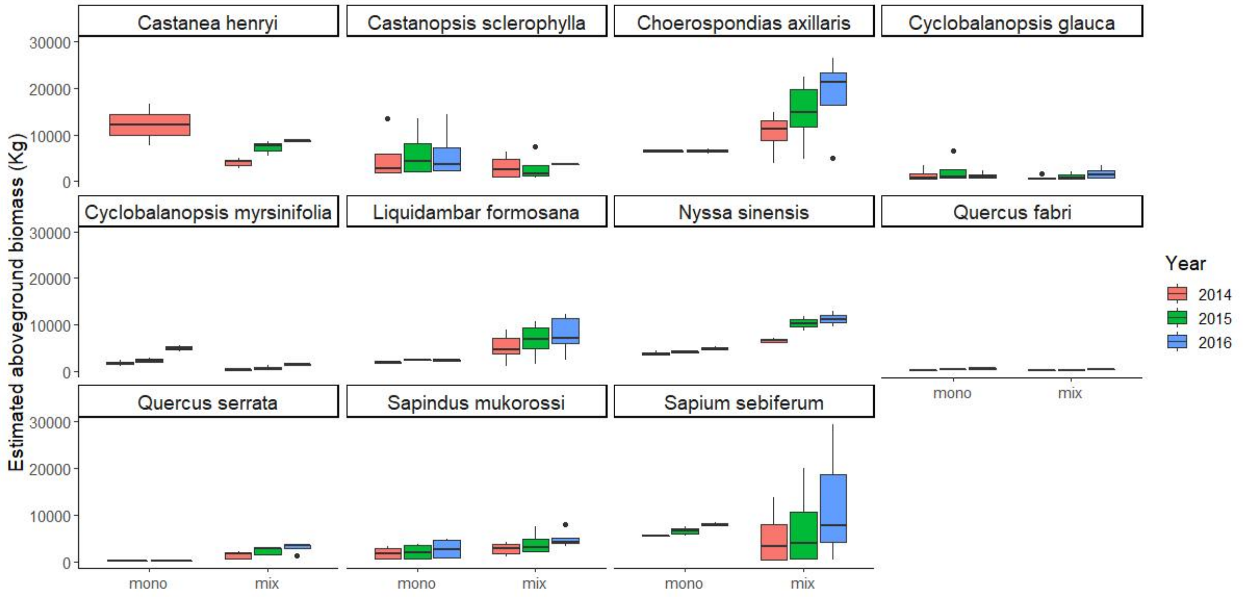


**Fig. S3.** Boxplot showing the estimated biomass of the trees from BEF China by species
